# Targeting the Long Non-Coding RNA LINC00599-205 splicing by novel candidate drug ABX464 to produce the anti-inflammatory microRNA miR-124

**DOI:** 10.1101/2019.12.11.871335

**Authors:** Laurent Manchon, Audrey Vautrin, Jamal Tazi, Aude Garcel, Noelie Campos

**Affiliations:** IGMM Université Montpellier, CNRS Montpellier, France; ABIVAX Therapeutics R&D department Montpellier, France

**Keywords:** long non-coding RNAs, splicing, regulatory, transcriptome assembly, capture sequencing, microRNA, anti-HIV, anti-inflammatory

## Abstract

Many nascent long non-coding RNAs have received considerable attention in recent years because of their major regulatory roles in gene expression and signaling pathways at various levels. Indeed long non-coding RNAs undergo the same maturation steps as pre-mRNAs of proteincoding genes, but they are less efficiently spliced and polyadenylated in comparison to them. Here we focus on a specific human long non-coding RNA and we show the activity of a new candidate drug that potentially affect its splicing and generate an anti-HIV and anti-inflammatory effects driven by upregulation of microRNA biogenesis. To investigate this activity we combine the use of capture sequencing technology and an *ab initio* transcript assembly on cells from six healthy individuals. The sequencing depth of capture sequencing permitted us to assemble transcripts exhibiting a complex array of splicing patterns. In essence, we revealed that splicing of the long non-coding RNA is activated by the drug whereas this splicing was not present in untreated samples.

## I. Introduction

Innovative application of new technologies in research is one of the major factors driving advances in knowledge acquisition. The combined use of capturing a targeted panel of RNA followed by next-generation sequencing (NGS) apparently suit this aspect of innovative study.

In this study, we conducted such combined approach to elucidate the mechanism of action of new anti-HIV ABX464 to better understand the means of the anti-inflammatory response in clinical trials. We highlight that ABX464 has no adverse effect on pre-mRNA splicing of cellular genes.

The tightly regulated process of precursor messenger RNA splicing is a key mechanism in the regulation of gene expression. Here, we reveal that ABX464 enhanced the expression and splicing of a single long non-coding RNA [1] to generate the anti-inflammatory microRNA [2] identified as being miR-124 both ex vivo and in HIV patients.

This specific capacity of ABX464 may present a safe way to cure HIV, by eliminating viral reservoirs, as well as to treat inflammatory diseases.

## II. Background & Summary

RNA sequencing (RNA-Seq) technologies [3][4] can provide an opportunity to assemble a complete annotation of the transcriptome, with techniques of *ab initio* transcript assembly [5] capable of identifying novel transcripts and expanding catalog of genes and their expressed isoforms that arise.

To profile such rare transcriptional events and thereby assess the full depth of the transcriptome, we employed a targeted RNA capture and sequencing strategy, which for brevity we term RNA CaptureSeq [6].

During the NGS process, hundreds of millions of cDNA segments are sequenced at the same time, in cycles from one or both ends. Without selective enrichment, any genomic region has an equal chance of being sequenced. With targeted gene capture, the proportion of DNA fragments containing or near targeted regions is greatly increased.

Briefly, RNA CaptureSeq involves the construction of tiling arrays across genomic regions of interest, against which cDNAs are hybridized, eluted and sequenced. Any region of interest in the genome can be targeted for enrichment by design, also our interest has focused on using targeted regions surrounding genomic loci of microRNA miR-124 for capture. Although this ability to isolate and target RNA has been used in genetic analysis for some time, here we combine this ability with deep-sequencing technology to provide an afforded coverage and permit the robust assembly to then discover novel mechanisms of transcriptional regulation [7].

As described in our related manuscript submitted earlier in Scientific Reports [8], our goal in this study is to elucidate the mechanism of action of ABX464, a new drug for curing HIV and treating inflammatory diseases. ABX464 is a small molecule that binds to the cap binding complex (CBC) [9], a complex at the 5’end of the pre-mRNA transcript that promotes the initial interaction with transcription and processing machinery [10][11]. According to that, we planned two experiences to obtain satisfactory results. To show that ABX464 does not affect cellular splicing we first used high-throughput RNA-Seq on peripheral blood mononuclear cells (PBMCs) [12][13], more concisely we used purified CD4+ T cells from the PBMCs of 4 donors. Additionally, to address the potential role of ABX464 in enhancing the expression and splicing of a single long non-coding RNA to generate the antiinflammatory miR-124 we used RNA CaptureSeq approach. Through this technic we also assessed the capability of ABX464 to generate spliced HIV viral RNA.

## III. Methods

### A. ABX464 does not affect cellular splicing

The CD4+ T cells were uninfected or infected with the YU-2 strain and were untreated or treated for 6 days with ABX464, followed by high-throughput RNA-seq. Each raw dataset of the samples contained between 44 and 105 million single-end reads (50 bp), with an average of approximately 60 million raw reads per sample (Fig. 1).

**Fig. 1.**
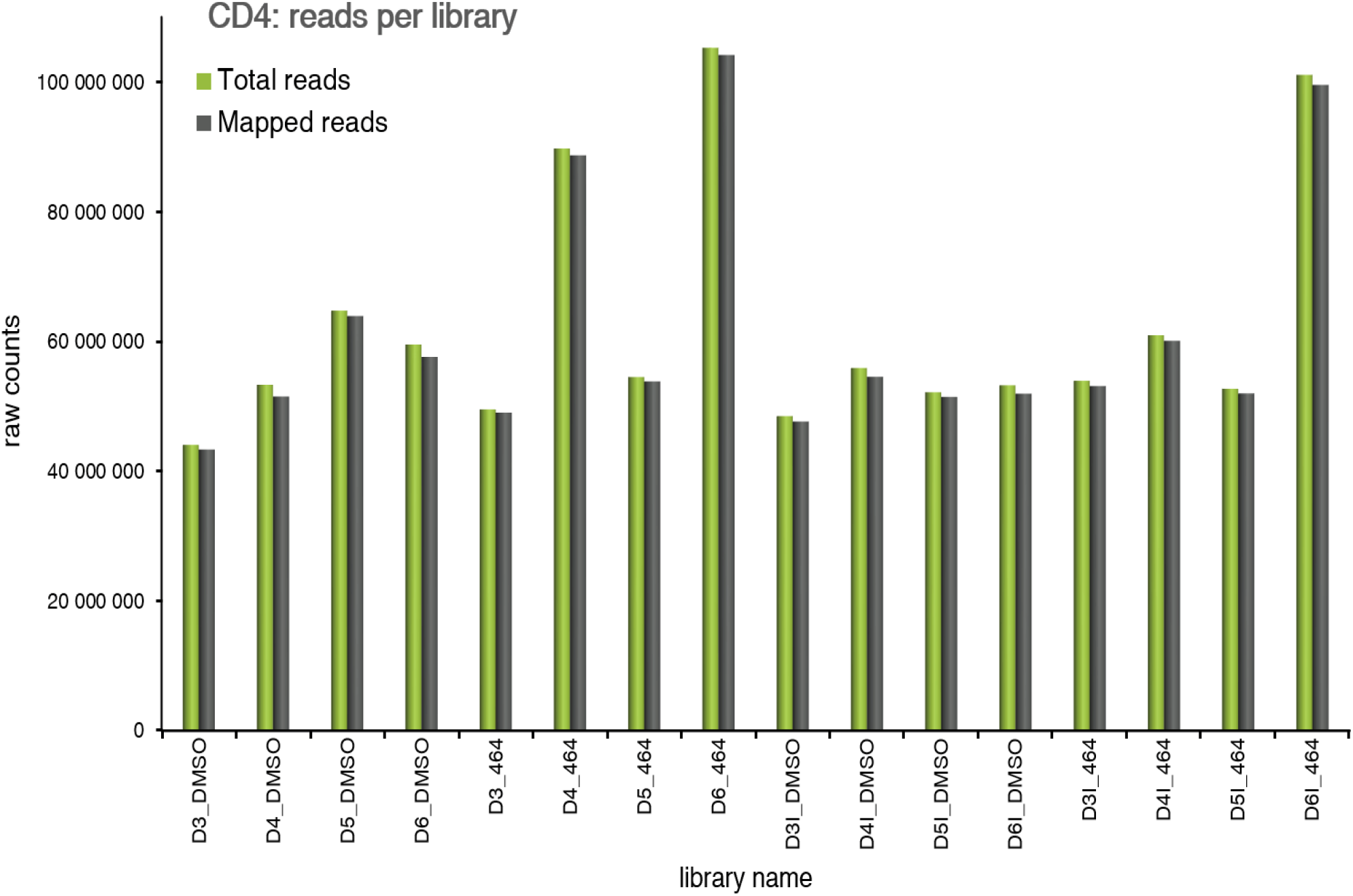
Read counts per RNA-seq librarie.

More than 97% of the bases had a quality score of ≥ Q20. Approximately 98% of the total raw reads were mapped to the human genome sequence (GRCh38), giving an average of 60 million human reads per sample for further analyses. The reads that were correctly mapped (approximately 98% of total input reads) to the gene and transcript locations (GTF annotation file) were subsequently analyzed using in-house package suites for transcript abundance normalization and evaluation.

For a gross estimation of the alternate splicing events (Fig. 2) modulated by ABX464 in CD4+ T cells, we compared junction read counts that originated from exon-exon boundaries across different samples. While the alignment of reads to exon junctions was purely coincidental, we did not observe gross differences in the total number of reads corresponding to exon-exon boundaries between treated and untreated CD4+ T cells, whether infected or uninfected, across any of the samples (Fig. 3).

**Fig. 2.**
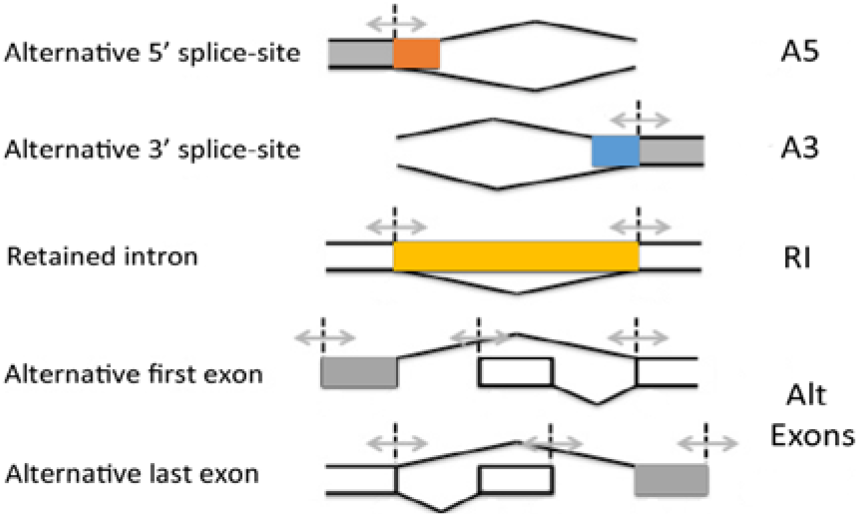
Classification of Alternative Splicing (AS) events of cellular genes

**Fig. 3.**
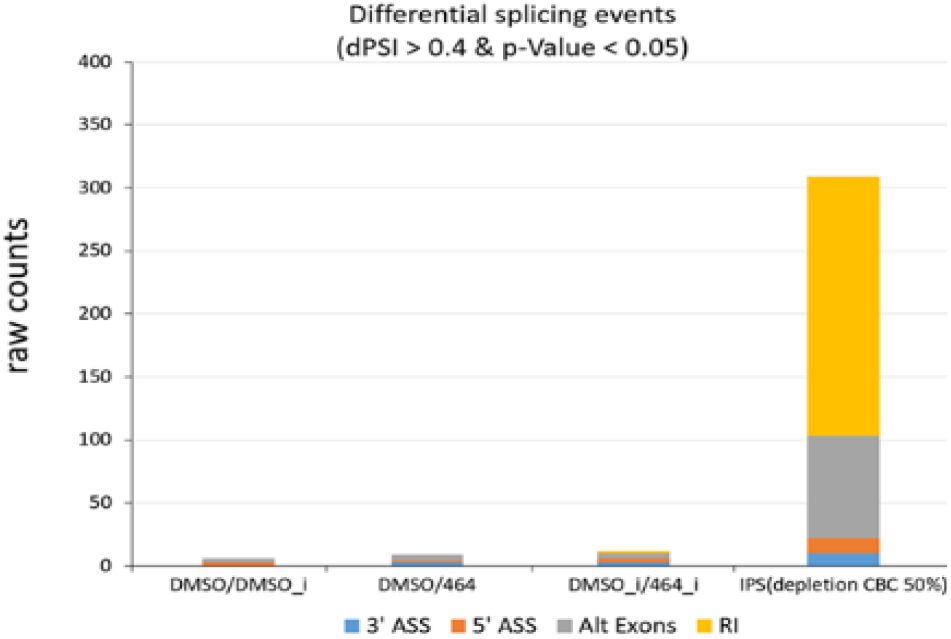
AS event counts comparing infected vs uninfected samples (DMSO_I vs DMSO_NI), uninfected vs uninfected treated by ABX464 (DMSO_NI vs 464_NI), infected vs infected treated by ABX464 (DMSO_I vs 464_I) and after 50% depletion CBC in IPS (IPS depletion of CBC by 50%).

For a more statistically qualified assessment of alternate splicing in these CD4+ T cells, we followed a method for differential splicing analysis across multiple conditions named SUPPA (super-fast pipeline for alternative splicing analysis). The alternate splicing events were classified into five major groups: Alternative 5′ splice site (A5SS), Alternative 3′ splice site (A3SS), Alternative first and last exons (Alt exons) and Retained exon (RI) (Fig. 3). A “percent splicing index (psi)” score (ψ-score) was calculated for each of the transcript variants for each sample. Using a stringent cutoff, the difference of ψ-score from untreated sample was fixed at 0.4 and p-value was fixed at 0.05 for differential splicing events induced upon treatment with ABX464 to be considered significant. We next calculated the ψ-scores of transcripts in uninfected versus infected CD4+ T cells (Fig. 3). No switchlike events were detected in infected or uninfected T CD4+ samples when compared to ABX464-treated samples (Fig. 3).

The exact numbers of common and differential splicing events between infected and uninfected CD4+ T cells treated with ABX464 were very low (fewer than 10 events), while depletion of NCBP1 (Nuclear Cap Binding protein subunit 1), a component of the CBC, in stem cells by only 50% gave rise to large variations, with 81 Alt exon, 12 A5’ SS, 10 A3’ SS and 206 IR events. The high level of IR indicated that CBC complex is a major component preventing accumulation of unspliced RNA as most of unspliced transcripts will be degraded by nonsense mediated decay (NMD). Comparison of exon coverage reads of a common highly expressed gene, B2M, between the ABX464 and DMSO conditions in the 4 donors revealed that ABX464 did not increase splicing events in B2M (Fig. 4).

**Fig. 4.**
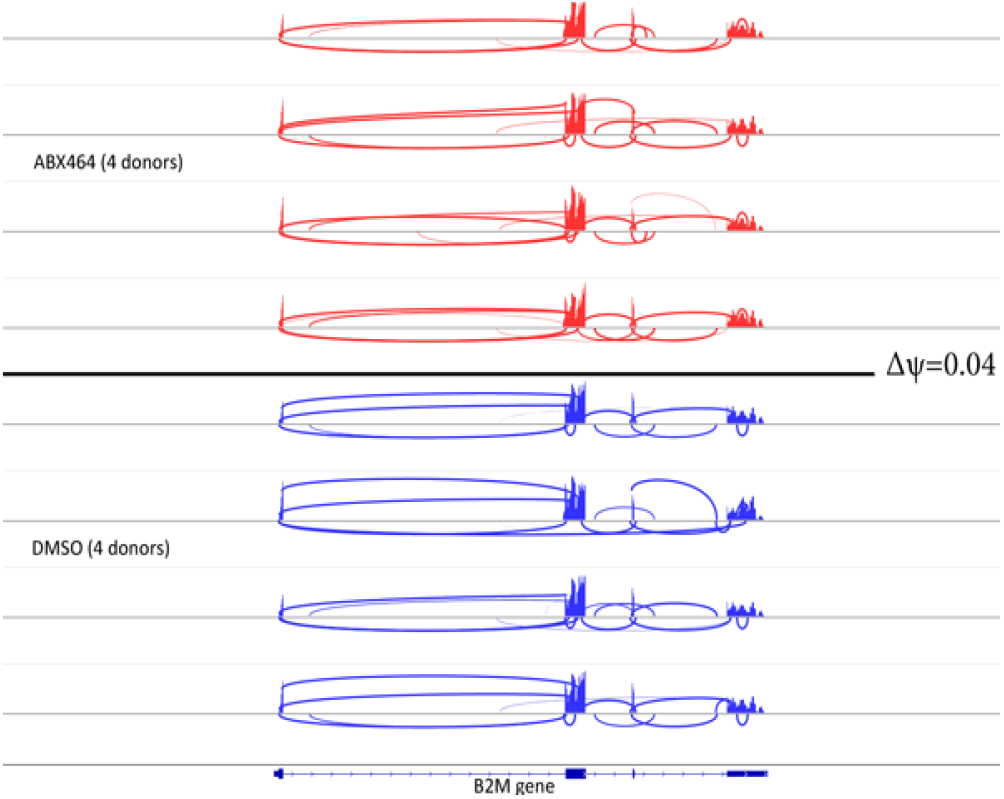
Comparing exon coverage reads of a common highly expressed gene (B2M) between the ABX464 and DMSO conditions in the 4 donors.

In summary, ABX464 treatment did not induce alternate splicing of transcripts, and therefore, ABX464 had no potential to dramatically alter gene expression in activated CD4+ T cells. Consistent with this result, FACS analysis of purified activated CD4+ T cells or PBMCs from 7 donors after 6 days of ABX464 treatment did not reveal any changes in the CCR6/CXCR3 or CD45/CCR7 (Th1 and effector memory cells, respectively) subpopulation.

To determine whether ABX464 induced a quantitative change in the transcriptome of infected cells, we measured transcript levels from the RNA-seq data. More than 60 000 different transcripts were detected among all samples. After counts per million normalization (CPM), transcripts of each sample were filtered out with a coverage cutoff value of CPM >5; that is, a gene was considered to be expressed in a sample if it had at least five counts for each million mapped reads in that sample. Since this experiment used four donors, four replicates were available for each condition (infected/uninfected cells with/without ABX464 treatment), and a gene was considered to be expressed if it was expressed to at least five counts for each million mapped reads in all replicates.

Following these selection criteria, we ended up with a list of transcripts common to all samples, including 11 700 different transcripts with an average of around 3000 raw counts per transcript. Using the CPM values of the qualified genes, a global expression plot was produced for each donor. Under the conditions of mild infection used in our study, we did not detect strong changes in gene expression, with only 15 genes being downregulated in response to infection in untreated samples (Fig. 5 top panel). ABX464 treatment resulted in the upregulation of 9 genes in the infected samples, and 6 downregulated and 7 upregulated genes in the uninfected samples (Fig. 5 middle and lower panels, respectively), demonstrating a mild effect of ABX464 treatment on gene expression.

**Fig. 5.**
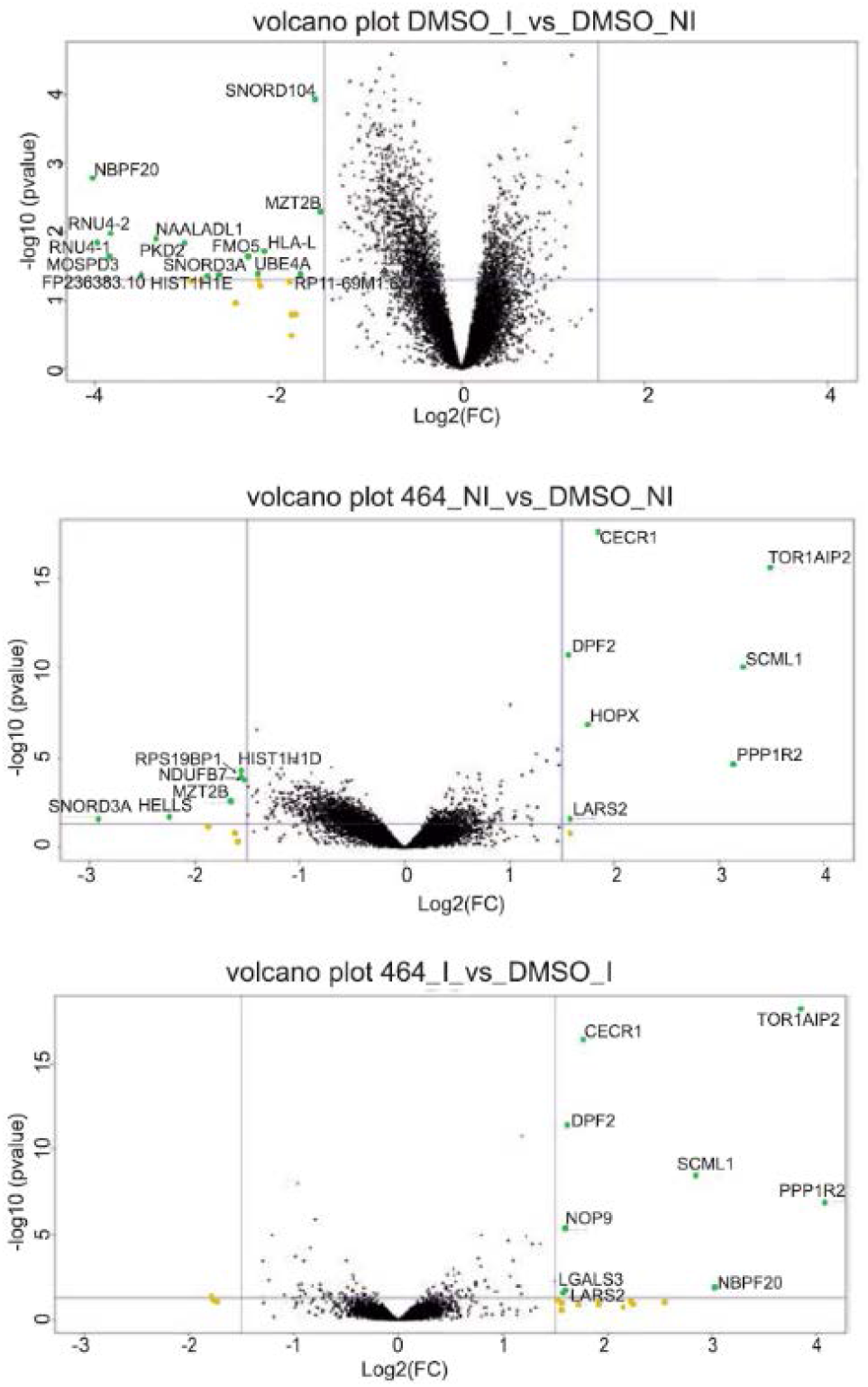
Volcano plot of DMSO_I vs DMSO_NI (upper panel), DMSO_NI vs 464_NI (middle panel), DMSO_I vs 464_I (lower panel). The gene expression variation generated by ABX464 treatment was very low in infected (6 downregulated genes) and uninfected (6 downregulated genes) samples.

### B. LncRNA splicing and upregulation of miR-124

Since ABX464 treatment did not induce any variation of the microRNA other than miR-124 (previously analyzed by microarray analysis on 104 microRNAs — data not shown) and very little variation of expression of genes that might host microRNA [2][14], we believe that ABX464-CBC interaction induces a specific effect of miR-124 biogenesis. This latter point is also demonstrated by the fact that miR-124 is transcribed by 3 loci but only the miR-124-1 locus is affected by ABX464 treatment and also the splicing of lncRNA LINC00599-205 is required for the production of miR-124-1. So, to directly assess the splicing of lncRNA known as LINC00599-205 RNA CaptureSeq together with *ab initio* transcript assembly was employed. The selected and targeted human DNA regions in this probe design were microRNAs and more specifically the miR-124 loci. Notably, miR-124 is represented in three different genomic loci [miR-124-1 (8p23.1), miR-124-2 (8q12.3) and miR-124-3 (20q13.33)]. We also used the miR-429 locus (1p36.33) as a control to evaluate its expression relative to miR-124. First approach was to evaluate expression level of each type of microRNAs (miR-124-1/-2/-3). We used Bowtie2 [15] in ‘very-sensitive’ mode to map the paired reads to chromosome 1, 8 and 20. We computed the mapped read counts using SAMtools [16] on 10-kb windows (5-kb surrounding each targeted locus). Raw counts of each type of microRNAs in conditions ABX464 versus DMSO were then reported in a table (Table 1).

**TABLE I.**

| LIB names | Raw Counts |  |  |  |
| --- | --- | --- | --- | --- |
|  | *Mir124-1* | *Mir124-2* | *Mir124-3* | *Mir429* |
| D1_464 | 116164 | 2580 | 21900 | 4098 |
| D4_464 | 51798 | 1204 | 5306 | 459 |
| D5_464 | 16352 | 1440 | 11709 | 2472 |
| D6_464 | 13294 | 5358 | 8498 | 6950 |
| D7_464 | 45480 | 17676 | 55708 | 7620 |
| D8_464 | 30950 | 8830 | 18253 | 5013 |
| D1_DMSO | 4014 | 704 | 6439 | 1760 |
| D4_DMSO | 7114 | 4 | 701 | 246 |
| D5_DMSO | 4748 | 4222 | 21941 | 3916 |
| D6_DMSO | 324 | 362 | 2542 | 678 |
| D7_DMSO | 2486 | 2326 | 14460 | 4109 |
| D8_DMSO | 1088 | 5452 | 7772 | 1482 |

Most of the aligned reads from the 10-kb regions of each locus originated from miR-124-1 and miR-124-3, whereas the number of aligned reads from miR-124-2 was quite low even after ABX464 treatment. The number of mapped reads from the three loci encoding miR-124 in the samples (Table 1) confirm that ABX464 led to a large increase in miR-124-1. In contrast, ABX464 had no effect on the expression of miR-429, located outside the miR-124 region, showing the specificity of ABX464 in targeting splicing of LncRNA.

The effect of ABX464 on the expression of miR-124 was most pronounced from the miR-124-1 locus. Close inspection of this region indicated that the miR-124 sequence is embedded in a long noncoding RNA (LINC00599-205) at the miR-124-1 locus. The miR-124-1 sequence is at the 3rd exon of lncRNA LINC00599-205, this is also a peculiar situation because most of microRNAs are generally classified into “intragenic” or “intergenic” miRNAs based upon their genomic location. Intergenic miRNAs, located between genes, are known to be transcribed as independent transcription units from their own Pol II or Pol III promoters, whereas intragenic miRNAs, embedded within introns or exons of genes on the same strand, are believed to be coregulated with their host genes by Pol II and hence share common regulatory mechanisms and expression patterns with their host genes [20].

Then to study the effect of ABX464 on splicing we studied the precursor mRNA (pre-mRNA) corresponding to miR-124-1. To do that we reconstructed all transcripts assembled within the precapture RNA-Seq. We did an assembly to build contigs from captured reads but selecting only the reads that map miR-124-1 locus.

Several assembly softwares exist and it was a tough decision point to choose one especially dedicated to assemble transcriptome, we decided to choose for this task Trinity software package [17] to piece the resulting sequence reads back together into contigs. Trinity, as other short read assemblers works in K-mer space, it has given us the best results and outperforms the others assembly programs not mentioned here. We generated 2 assemblies, one per condition: DMSO and ABX464 treatment. Due to the lack of reads in this small target region and for merely reasons we have pooled reads from all the donors to get best assembly. Afterwards, we used GMAP aligner [18] to align the contigs and visually explored the alignment results in IGV viewer [19] to select contigs of interest. Three tracks were displayed in IGV: ABX464, DMSO and the alignments read depth (read coverage) corresponding to ABX464 track (Fig. 6).

**Fig. 6.**
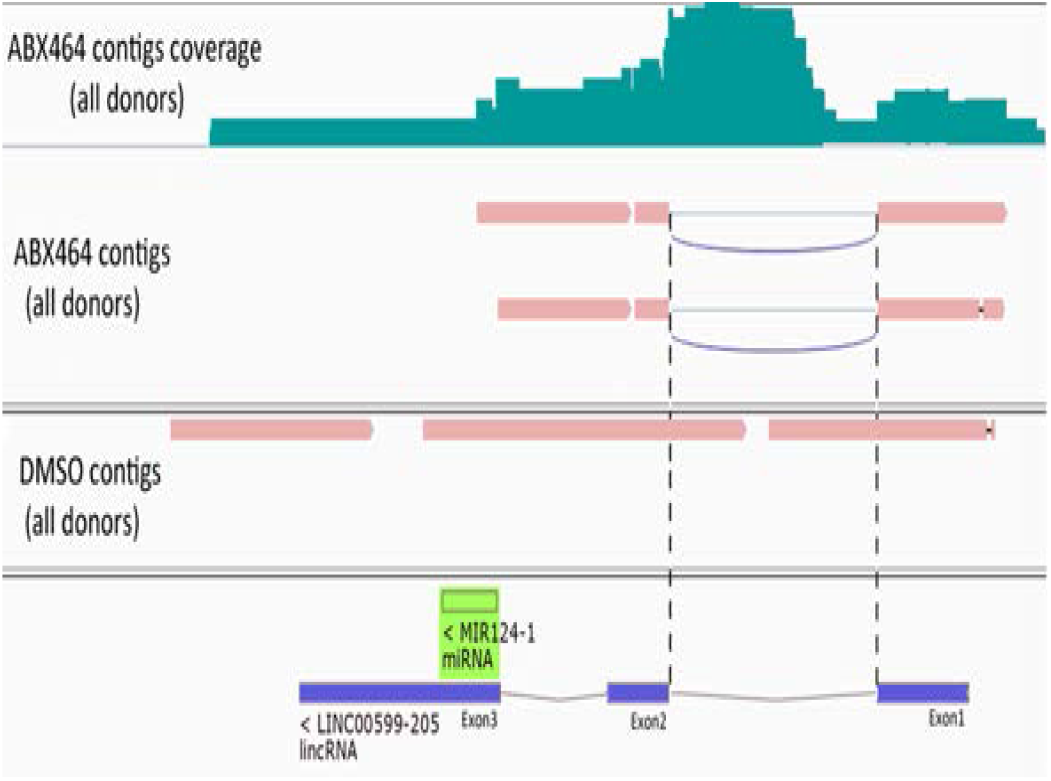
Locus of long noncoding RNA (LINC00599-205) whose splicing is stimulated by ABX464.

In addition, we have checked ‘Show junction track’ in IGV viewer to visualize splice junctions. Formerly we have obtained one splicing region in ABX464 condition that didn’t appear in DMSO condition and confirmed that ABX464 treatment induce splicing on pre-miR-124-1 RNA. To confirm this point of view we used SAMtools to count the reads mapped to exon/intron and exon/exon junctions localized inside pre-miR-124-1 locus then plotted the counts to produce a histogram (Fig. 7).

**Fig. 7.**
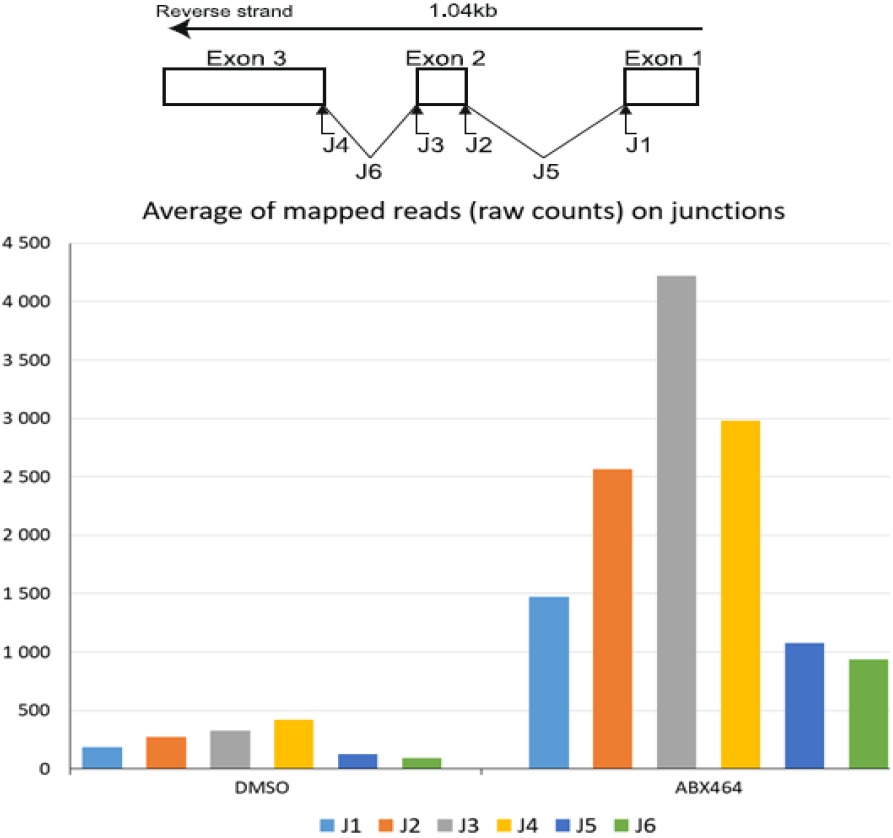
Raw counts of the reads at splice junctions (J1, J2, J3 and J4), exon-exon (J5 and J6) quantified by RNA CaptureSeq in PBMCs treated with ABX464 versus control DMSO.

This approach revealed that splicing of lncRNA LINC00599-205 is activated by ABX464, and this splicing was not present in untreated samples. Most of the contigs in the untreated samples aligned with unspliced lncRNA LINC00599-205. Since spliced RNA is more stable than unspliced RNA, we must consider that spliced lncRNA LINC00599-205 is a source of miR-124 upregulation and therefore becomes less stable.

To directly assess the contribution of lncRNA LINC00599-205 to the production of miR-124, we cloned the genomic sequence of chromosome 8 from 9 903 167 through 9 904 210 of lncRNA LINC00599-205 into a plasmid vector (Fig. 8A). Both the spliced and unspliced transcripts of lncRNA LINC00599-205 could be readily detected in the transfected HeLa cells (Fig. 8B). However, we failed to detect an upregulation of miR-124 in transfected HeLa cells in response to ABX464 treatment since splicing is maximized in HeLa cells and the CBC complex would have an effect at the site of transcription (Fig. 8B).

**Fig. 8.**
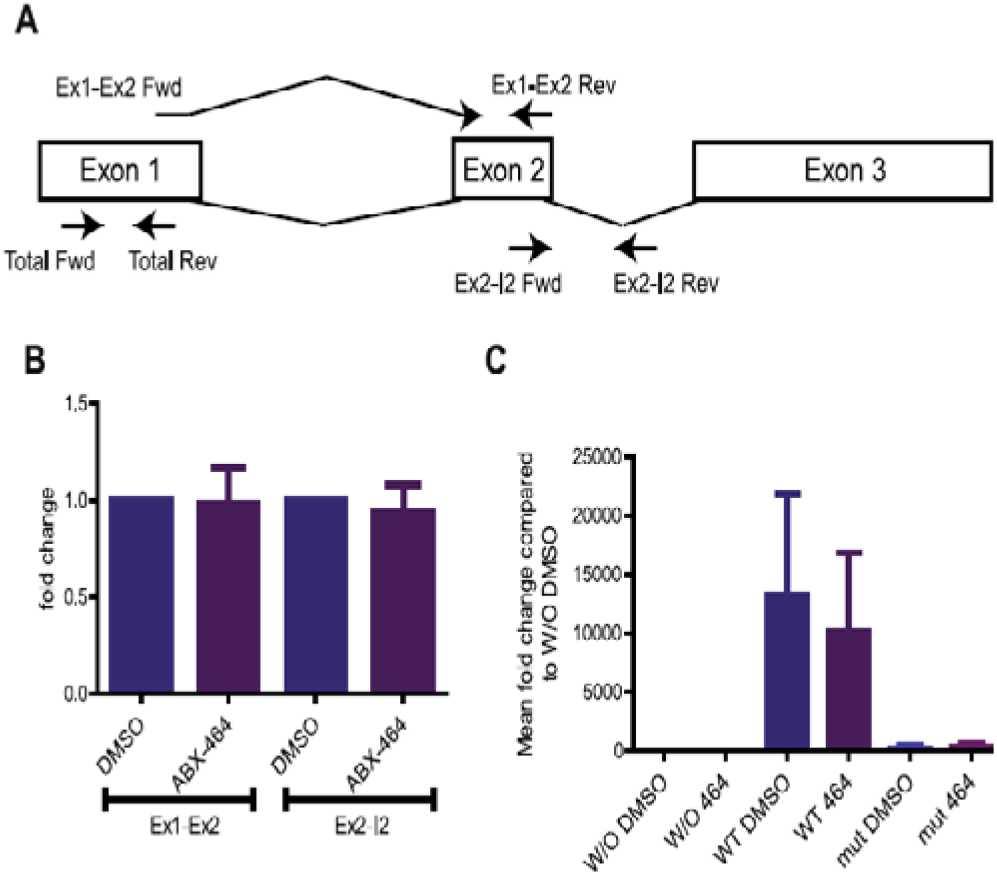
Splicing of LncRNA LINC00599-205 is required for the production of miR-124.

However, transient transfection of the plasmid into HeLa cells resulted in a dramatic (1 250-fold) increase in miR-124 levels (Fig. 8C, left panel). In contrast, when the splice sites of the lncRNA LINC00599-205 are mutated only traces of miR-124 are detected (Fig. 8C, left panel). Consistent with the fact that unpliced lncRNA LINC00599-205 are less stable, the amount of mutated lncRNA LINC00599-205 is less than wild type (Fig. 8C, right panel).

## IV. DISCUSSION

Our investigation provides new insights into the mechanism of action of ABX464 in upregulating the antiinflammatory miR-124 in patients. ABX464 binds CBC, a complex that stimulates processing reactions of capped RNAs, including their splicing, 3′-end formation, degradation, and transport. Given that the CBC is believed to bind all classes of m7G-capped RNAs, including precursors and mature forms of mRNAs, stable long noncoding RNAs (lncRNAs), nonadenylated histone RNAs, and precursors of spliceosomal small nuclear RNAs (snRNAs), it is important to understand how the binding of ABX464 to CBC complex achieves its specificity and robustly induces the variation of splicing of lncRNA LINC00599-205.

We don’t know the reason for that. It could be that weak splice sites that characterizes both transcripts, is at the origin of the induction of these splicing events by ABX464-CBC complex. Since unspliced lncRNA LINC00599-205 cannot exit the nucleus due to quality control machinery and CBC-mediated control, spliced lncRNA LINC00599-205 constitutes a miRNA storage form, possibly in addition to other functional properties of the intact spliced transcript. This storage may be maintained through low transcriptional and degradative activity of unspliced lncRNA LINC00599-205 and producing only low levels of mature miR-124 release under normal conditions.

A significant progress has been made regarding the role of lncRNAs and has expanded our understanding of how their regulation represent an important mechanism for altering changes in gene transcription. In this paper we highlight the splicing of lncRNA LINC00599-205 is important to stabilize the mature transcript for recognition by the microRNA processing machinery and thereby produce miR-124. Consistently, the splicing of lncRNA LINC00599-205 is a prerequisite for the production of miR-124. ABX464 treatment could, thus, enable the rapid release of a large amount of miR-124 through lncRNA LINC00599-205 splicing (Fig. 9).

**Fig. 9.**
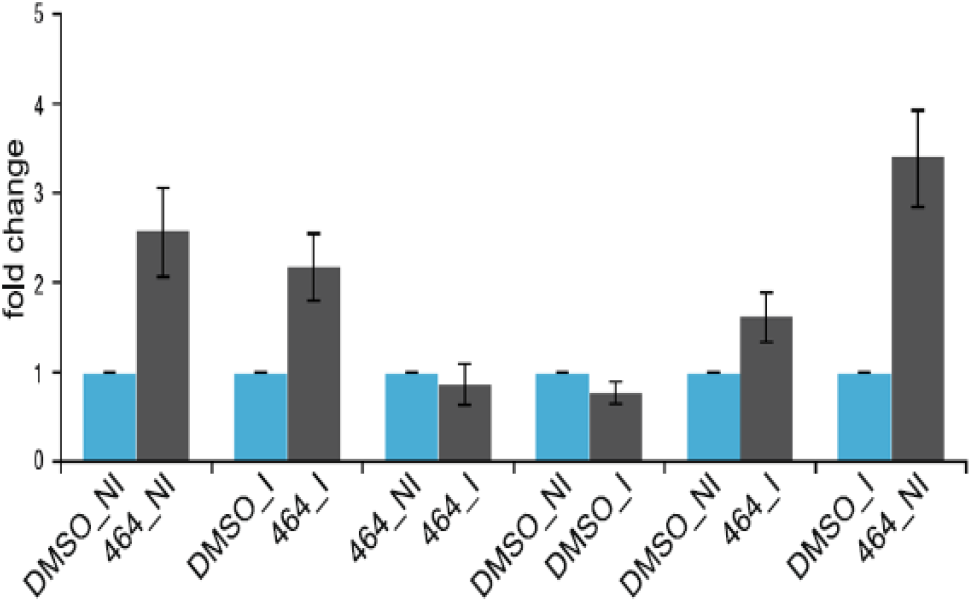
Cells Quantification of miR-124 expression using TaqMan Low Density Array technology in CD4+ T cells.

## ACKNOWLEDGMENT

We thank all members and all the contributors for their work to this paper. This work was supported by the collaborative laboratory ABIVAX-CNRS.

